# The evolution of democratic peace in animal societies

**DOI:** 10.1101/2023.02.14.528470

**Authors:** KL Hunt, M Patel, DP Croft, DW Franks, PA Green, FJ Thompson, RA Johnstone, MA Cant, DWE Sankey

**Affiliations:** Centre for Ecology and Conservation, Faculty of Environment, Science and Economy, University of Exeter, Penryn Campus, Cornwall TR10 9FE, UK; Centre of Excellence for Data Science, Artificial Intelligence and Modelling and Department of Biology, University of Hull, Hull, HU6 7RX, UK; Centre for Research in Animal Behaviour, Faculty of Health and Life Sciences, University of Exeter, Exeter, EX4 4QG, UK; Department of Biology and Department of Computer Science, University of York, York YO10 5DD, UK; Department of Ecology, Evolution, and Marine Biology, University of California, Santa Barbara, CA, 93106, USA; Department of Zoology, University of Cambridge, Cambridge, CB2 3EJ, UK

**Keywords:** Leadership, Intergroup conflict, Democracy, Game theory, Evolution

## Abstract

A major goal in evolutionary biology is to elucidate common principles that drive human and other animal societies to adopt either a warlike or peaceful nature. One proposed explanation for the variation in aggression between human societies is the ‘democratic peace’ hypothesis. According to this theory, autocracies are more warlike than democracies because autocratic leaders can pursue fights for private gain. However, autocratic and democratic decision-making processes are not unique to humans and are widely observed across a diverse range of non-human animal societies. We use evolutionary game theory to evaluate whether the logic of democratic peace may apply across taxa; specifically adapting the classic Hawk-Dove model to consider conflict decisions made by groups rather than individuals. We find support for the democratic peace hypothesis without mechanisms involving complex human institutions and discuss how these findings might be relevant to non-human animal societies. We suggest that the degree to which collective decisions are shared may explain variation in the intensity of intergroup conflict in nature.

## MAIN TEXT

The tendency to organise into groups and engage in collective violence against other groups is a striking and destructive feature of human behaviour ^1–3^. Such intergroup conflicts have persisted for many tens or hundreds of thousands of years, and continue take a tremendous toll on human life ^4–7^. Therefore, understanding the root causes of intergroup conflict is an issue of significant and urgent importance. Until recently, human intergroup conflict was seen as a particularly human phenomenon, predicated on uniquely human cultural traits ^2,3,8,9^. Now, however, our increasing understanding of intergroup conflict in other animals has helped to illuminate the evolutionary forces that may have shaped the phenomenon ^10,11^. Across taxa there is great variation in the severity and frequency of intergroup conflict ^12–14^, which provides an opportunity to identify shared traits which drive societies towards war or towards peace.

### Democratic peace in humans and other animals

In human societies, a dominant explanation for variation in interstate conflict is the democratic peace hypothesis, which proposes that democracies are less prone to initiate interstate conflicts than autocracies because shared decision-making acts as a restraint on warmongering (or ‘exploitative’) leaders ^8,13,15–20^. This hypothesis has received widespread empirical support ^21–24^. For example, a recent analysis of patterns of conflict among 186 countries over a 42-year period found the statistical link between democracy and peace to be five times stronger than that between smoking and lung cancer ^24^. Until now it has been assumed that democratic peace relies on uniquely human institutions ^25–27^. However, in principle the logic of democratic peace could apply to intergroup conflict among any biological groups that exhibit shared or collective decision making, suggesting that the hypothesis may have much broader scope across other taxa.

Here we use evolutionary game theory to explore the potential for democratic peace to explain variation in intergroup conflict in biological societies, particularly animal societies. A large body of literature now demonstrates that collective movement decisions in animal societies can be either unshared (i.e., dictated by leaders; ^28–34^) or shared across the group (i.e., democratic ^35–41^). These democratic decisions are observed when the majority can influence group movements more than any one individual or subset of individuals. Shared and unshared collective movement decisions could translate into collective decisions to initiate conflict too. For example, if a leader was motivated to fight another group, it could lead its group in a hostile move towards that rival’s territory. Equally, in a shared decision-making context, if the majority of a group prefer to avoid a fight, they may collectively form a consensus to retreat and evade the rival. Both intergroup conflict and collective movement decisions are observable across many taxa, from bacteria ^42,43^ to primates ^30,35,44^, and therefore collectively deciding whether to fight could be, in theory, achievable by a wide range of social organisms.

### The model

Given that many social organisms have the necessary prerequisites for democratic peace, we explore whether evolution would favour such a relationship in an evolutionary game theoretical model (based upon a Hawk Dove game ^45^) in which collectives rather than pairs compete over resources. In an infinite population, groups of size *N* encounter each other randomly, and collectively decide to play either a peaceful, Dove, or aggressive, Hawk, strategy. Upon meeting, two Doves will share a resource, *V*, equally. Whereas a Dove will concede the resource to a Hawk (Hawk = *V*, Dove = 0). The disadvantage to playing Hawk is that when two Hawks meet, they will fight, resulting in costly losses, *C*, for one group (and *C* > *V*), such that playing Dove can be advantageous. Within each group, individuals are divided into two distinct classes: leaders with proportion ε and followers with proportion 1-ε. The classes differ in their decision-making influence and the costs and benefits of intergroup interactions as described in the following two sections.

### Collective decisions

Both classes evolve strategies with probability of Hawk-playing *P*_*L*_ for leaders and *P*_*F*_ for followers. Dove is played with probability 1 – *P*_*L*_ by leaders and 1 – *P*_*F*_ by followers. These strategies are combined into a collective decision, whereby their entire group either play either Hawk or Dove with probability *P* (Eqn. 1). However, all votes are not necessarily counted equally. The leaders’ probability for playing Hawk always contributes toward the group’s collective decision, such that the leaders’ combined influence on the group strategy, *P*, is proportional to *P*_*L*_ε. Follower strategies also have an influence on *P*, except that it is weighted by a shared decision-making parameter Ω, such that the followers’ combined influence is proportional to *P*_*F*_ Ω(1-ε) where (0 ≤ Ω ≤ 1). When Ω = 0, follower strategies are weighted by zero, so only the leaders have influence, and the decision-making structure can be described as an unshared consensus decision ^51^. In contrast, when Ω = 1, any given follower and any given leader both have equally shared influence, and the decision-making structure can be described as a shared consensus decision ^51^. Note, our model assumes that once a collective decision is made, the whole group abide by the decision. Thus, applicable to societies where the possible benefits of defection are outweighed by the costs of punishment^46,47^, isolation from the group^48,49^, or delays in decision-making^50^.

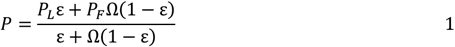

### Distribution of costs and benefits

Leaders and followers are further differentiated by receiving different shares of the costs, *C*, and benefits, *V*, from fighting, governed by the parameters *dc* and *dv* (where 0 ≤ *dc* ≤ 1 and 0 ≤ *dv* ≤ 1; see Equations 2-5 in Materials and Methods). When *dc* = 0.5, the costs paid by leaders, *C*_*L*_, and followers, *C*_*F*_, are equal. But when *dc* > 0.5, leaders receive an advantage by avoiding the costs of fighting relative to followers (*C*_*L*_ < *C*_*F*_) (Fig. S1). This could be because leaders are larger, tougher, better armoured, or able to take up safer positions during the fight such as at the back of the group ^7,13,20^. Likewise, when *dv* > 0.5 leaders’ share of the benefit from the antagonistic interaction, *V*_*L*_, is greater than that of the groups’ followers, *V*_*F*_. This represents cases where leaders are socially dominant to followers and able to displace them from contested resources such as food or mates that obtained as a reward after victory ^29,30,52,53^. We also consider cases where followers pay lower costs, dc < 0.5, or gain more reward, dv < 0.5, following the literature suggesting that there can be greater costs associated with leadership ^54–56^. We can use the values of *C*_*L*_, *C*_*F*_, *V*_*L*_ and *V*_*F*_ to calculate the probability of hawk-playing that maximises the fitness of each player class. For leaders this optimum strategy is equal to *V*_*L*_ / *C*_*L*_, whereas for followers it is equal to *V*_*F*_ / *C*_*F*_.

**Table 1.** Parameter table.

| *Parameter or variable* | *Description* |
| --- | --- |
| $\Omega$ | Shared decision parameter – the degree to which collective decisions are shared from 0 = complete control by leaders to 1 = equal share of decisions by leaders and followers |
| $\epsilon$ | Proportion of leaders in each group. Proportion of followers = 1 - $\epsilon$ |
| $N$ | Number of individuals in each group |
| $V$ | The value of the contested resource |
| $C$ | The cost of losing a conflict |
| $dv$ | Distribution of $V$ across the classes. Higher values indicate leaders monopolising the resources (Fig. S1) |
| $dc$ | Distribution of $C$ across the classes. Higher values benefit leaders, in that followers pay the majority of the costs (Fig. S1) |

### Virtual evolution of leader and follower strategies

We initialise a model where both leaders and followers start with a majority Dove strategy (*P*_*L*_ = *P*_*F*_ = *0*.*01*). We then proceed to allow leader and follower strategies to update in an iterative process until both classes stabilise on an evolutionarily stable strategy. In each timestep we calculate the relative fitness of playing either Hawk (*W*_*LH*_ and *W*_*FH*_ for leaders and followers respectively) or Dove (*W*_*LD*_ and *W*_*FD*_ respectively) (Eqns. 6-9 in Materials and Methods). If either class were found to perform better playing Hawk (for leaders *W*_*LH*_ > *W*_*LD*_, or for followers *W*_*FH*_ > *W*_*FD*_), then their class evolves to become more aggressive in the next timestep. Vice versa, if playing Hawk performs worse than Dove, then strategies become more peaceful. This iterative process continues until a stable strategy has been found. Each class’s strategy influences the others’ (through changes to *P*; Eqn. 1), but both update their hawk-playing probabilities (*P*_*L*_ *and P*_*F*_) independently in each timestep, allowing the model to find a different stable strategy for each class.

### The evolution of democratic peace (or democratic war)

We find that when the leaders are advantaged – meaning that on average they pay less of the costs of fighting (*dc >* 0.5) or gain more of the rewards (*dv* > 0.5) relative to their followers – they exhibit a higher probability of Hawk playing than the followers, who play an overall more peaceful strategy (Fig. 1; Fig. S2). In these scenarios, our model indicates support for the core prediction of the democratic peace hypothesis, in that increasing the follower’s share of the collective decision, Ω, results in a decrease in the aggressive Hawk-playing strategy of the population *P*.

**Figure 1:**
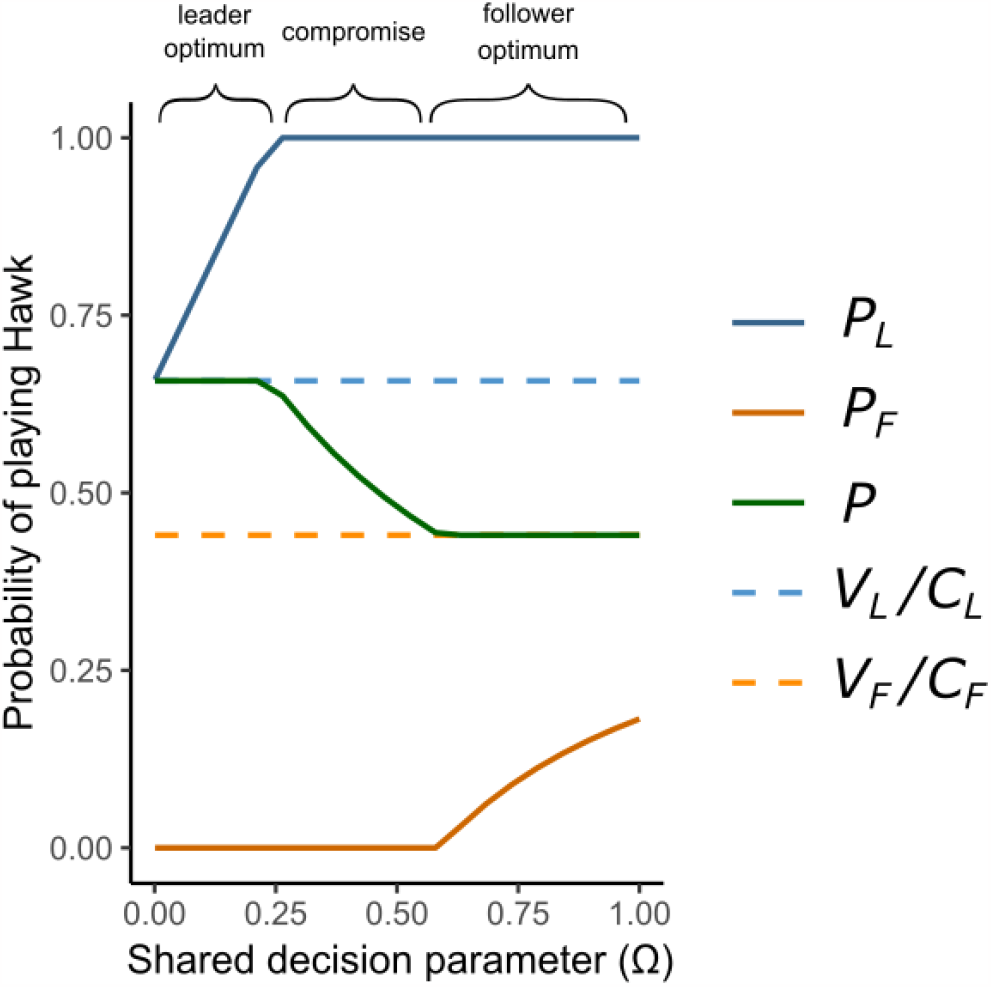
The evolution of democratic peace. The strategy of Hawk-playing for leaders (*P*_*L*_) and followers (*P*_*F*_) and the probability of playing Hawk across the population (*P*) are shown against shared decision parameter (Ω). ***Leader Optimum***. When leaders wield full control over the group’s decision-making (Ω = 0) the group’s probability of playing Hawk (*P*) is equivalent to the leaders’ optimum probability (*V*_*L*_ / *C*_*L*_). As Ω increases above 0, the follower’s strategy (*P*_*F*_) begins to have increased influence on the group’s probability of playing Hawk (*P*). Leader strategies respond to compensate for this increased follower influence by increasing their hawkishness (*P*_*L*_), which acts as an anchor and ensure that the group’s strategy (*P*) does not deviate from the leader’s optimum. This trend continues with increasing values of Ω until the leaders become obligate Hawk players (*P*_*L*_ = 1), at which point they cannot increase their strategy to become any more aggressive. ***Compromise***. Once leaders have become obligate Hawk players the group’s probability of Hawk-playing (*P*) begins to decrease. The followers increasing influence works to sway the group’s strategy (*P*) away from the leader’s preference (*V*_*L*_ / *C*_*L*_) and towards the follower’s preference (*V*_*F*_ / *C*_*F*_). In compromise states, both classes are observed playing strategies of either obligate Hawk or obligate Dove, for leaders or followers respectively. ***Follower optimum***. At higher values of shared decision-making parameter Ω, followers will have sufficient influence over the decision-making process to ensure that the group’s played strategy (*P*) matches their optimum (*V*_*F*_ / *C*_*F*_). For the highest values of Ω the followers respond by increasing their likelihood of playing Hawk (*P*_*F*_). This adjustment is required to account for the diminishing relative influence of the leader’s strategy (*P*_*L*_) on the group’s played strategy (*P*), and to keep the group-level outcome in line with their fitness optimum (*V*_*F*_ / *C*_*F*_). Parameter values: ε = 0.3, *C* = 2*V, dc* = 0.55, *dv* = 0.55.

This assumption that the leaders generally benefit more than the followers is well supported in nature, because leadership is often associated with older, dominant, or otherwise privileged, high-status individuals who can benefit from priority of access to resources, including those that are gained from fighting ^29,30^. Similarly, leaders may be able to lessen their individual costs of fighting relative to followers on account of being larger, stronger, or being able to occupy safer positions during the fight ^7,13,20^. However, if these assumptions are not met and leadership is instead costly and disadvantageous relative to being a follower ^53,54,56,57^, then increasing shared decision-making can instead increase the hawkishness within the group in a phenomenon we describe as “democratic war” (Fig. S3). A key insight from our model is that we find that the democratic peace is only upheld when leaders are advantaged relative to followers, otherwise democracy has the opposite effect in increasing aggression in intergroup interactions.

To better understand whether democratic peace or war is more likely in a given biological system, it is useful to attempt to quantify the inter-individual distribution of the costs and benefits. This requires measuring and quantifying our *dc* and *dv* parameters. Empiricists could achieve this by measuring proxies for the costs of fighting, such as mortalities or injuries, and examining whether these are distributed evenly among group members or are biased either towards or against individuals with more decision-making responsibility. Similarly, empiricists could measure how resources, such as food or reproductive opportunities, are shared after the conflict to approximate the division rules used to share the benefits of fighting. An example of estimating these parameters in practice comes from banded mongooses, where researchers observed that females, who often act as leaders, are disproportionately less likely to die from intergroup fighting (high *dc*) and are also more likely to gain from mating opportunities (high *dv*) ^13^. Given the females have both greater decision-making influence and are advantaged (Fig. S2), the high levels of violent conflict observed in this system are consistent with the predictions from our model (and that of ^13^).

### Loudest voice prevails

Perhaps surprisingly, when followers have sufficient influence, they can control the collective decision completely in their favour, not because each follower has more influence than each leader, but because followers are in the majority. When ε < 0.5 (as in Fig. 1) the combined influence of followers can outweigh the leaders’ strategy. Intuitively, the strategy of leaders can be overturned by smaller values of follower influence Ω when the proportion of leaders ε is small but require much larger values of shared decision-making parameter Ω, or may not be overturned at all, when the proportion of leaders ε is large (Fig. 2; also see Fig. S4 for a broader parameter space).

**Figure 2:**
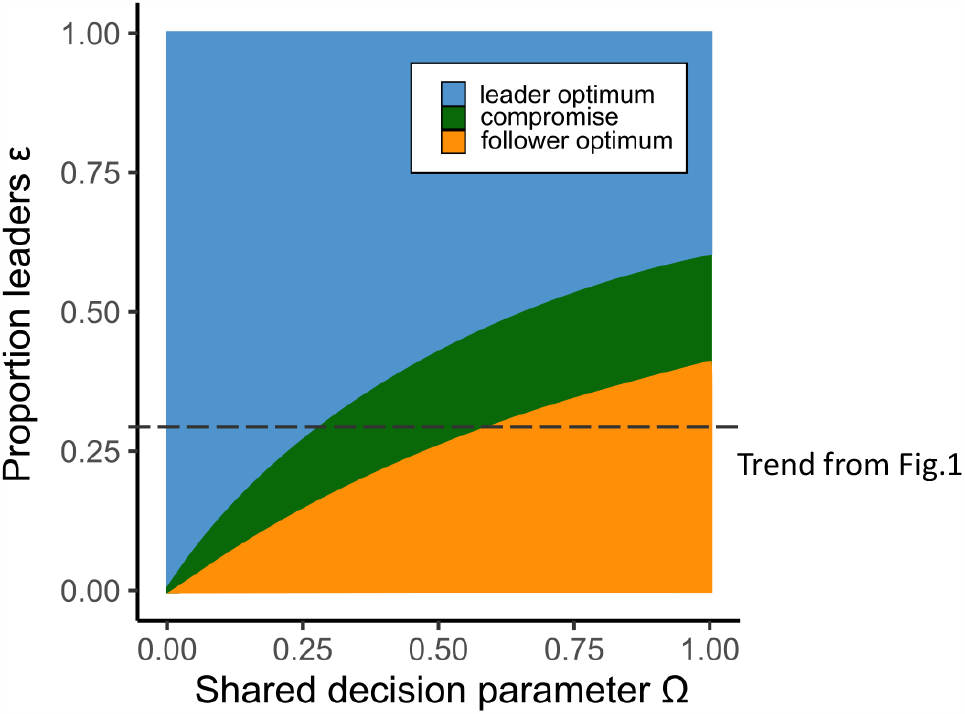
Follower control of decision is only possible when they are the majority class and decisions are at least partially shared. Population level outcome for the entire range of possible shared decision parameter values (Ω) and leader proportions (ε). Outcome given as the follower’s optimum (*P* = *V*_*F*_ / *C*_*F*_), leader’s optimum (*P* = *V*_*L*_ / *C*_*L*_), or compromise (*V*_*F*_ / *C*_*F*_ < *P* < *V*_*L*_ / *C*_*L*_). Dashed line indicates the same parameter space displayed in Fig. 1. Parameter values *C* = 2*V, dc* = 0.55, *dv* = 0.55.

There is a conspicuous absence of compromise at both low and high levels of democracy with an outcome in favour of either leader or followers respectively. This emerges because of a “loudest voice prevails” dynamic ^58,59^, in which both classes adopt obligate Hawk or Dove across much of the parameter space, yet one party is unable to prevent the other from achieving its optimum outcome. The loudest voice prevails outcome is similar to the predicted resolution in other evolutionary conflicts. In models of genomic imprinting, paternal and maternal genes may evolve all-or-nothing expression to either be active or completely silenced ^58,59^. In eusocial Hymenoptera, colonies are observed to produce a majority of males or females ^60^, which has the effect of driving the population ratio of males to females towards their optima ^61^. Further parallels can be found in humans, where political party members are known to elect leaders with more extreme views than themselves to negotiate with rivals because extreme views act as better negotiation anchors and more often succeed in passing moderate legislation that is closer to the party member’s original preference ^62^.

The success of extreme strategies in ‘anchoring’ and controlling the collective decision in our model is determined by the difference between the extreme strategy (*P* = 1 for Hawk or *P* = 0 for Dove) and the optimum of a given class. When this difference is low – i.e., the ‘extreme’ strategy closely resembles the optimum – then that player class will be less successful in controlling the decision outcome in their favour. For example, if a class’s optimum probability of Hawk playing was equal to 0.2, an extreme strategy for Dove (*P*_*L*_ or *P*_*F*_ = 0) would only have limited impact, whereas an extreme strategy for Hawk (*P*_*L*_ or *P*_*F*_ = 1) would be more effective at anchoring and controlling the collective decision. Importantly, in our model strategies are bounded between 1 and 0, as a group cannot play Hawk or Dove more than 100% of the time. This represents an important distinction between models of collective decision-making with bounded actions (as here) to models with continuous actions (e.g., decisions over the time to depart a foraging patch – as in ^63^).

### Interpretation for empirical systems

In our model, the empirical interpretation of the shared decision-making parameter Ω is intentionally left unspecified. This is because we imagine different possible ways in which Ω might vary both within, and between, species. Firstly, we propose that Ω could be a species-specific parameter, in that it describes the distribution of decision-making influence within a given animal society. In line with this, many recent studies and comparative analyses have described the variation in decision-making between different taxa as being more shared ^29,35–41^, or unshared ^28–34^, which would correspond to high and low values of Ω respectively. For example, olive baboons (*Papio anubis*) make shared consensus decisions when deciding where to travel as a group (Ω = 1) ^35^, whereas bottlenose dolphins (*Tursiops sp*.) make unshared consensus decisions where males have disproportionate influence relative to the rest of the group (Ω < 1)^31^. It is important to note that much of the research in collective decision-making in animals has thus far focused on group movement and foraging decisions ^30,33,36,37,39,64^. It is largely unknown to what extent the same decision-making rules, and therefore Ω values, are generalisable across different contexts (but see ^29^). For example, meerkats (*Suricata suricatta*) initially share the decision of when to stop foraging ^39^, but dominant females then dictate the decision of which burrow they go home to ^34^. Leadership dynamics in chimpanzees *(Pan troglodytes*) are similarly fluid with different individuals wielding influence depending on the context of group movement, within-group conflict resolution or between group aggression ^65–67^. As our model is focused on intergroup conflict, it is important that future work considers the distribution of decision-making (Ω) within an intergroup conflict context, which may not be the same as during other contexts (e.g., group foraging ^29,68^).

Secondly, we consider the possibility that Ω might vary within social groups of the same species, based on the group’s unique composition of individuals. Leadership is often governed by the phenotypes of individuals, such as their age ^29,32,33,44,64^, sex ^33,44,64,68–71^, or personality ^72–74^, and therefore the distribution of decision-making (Ω) within a social group may vary over time with changes to group composition. For example, in a chimpanzee troop, their propensity to engage in collective hunts declined after the death of an “impact hunter” ^75,76^, which may reflect resultant changes to the decision-making dynamics within the group. Although group hunting is different to intergroup conflict, there are similarities between the two behaviours ^77,78^, and this example demonstrates how demographic changes can influence collective decision-making and ultimately group-level behaviours. Demographic changes might have the most impact on the decision-making parameter (Ω) when they involve the presence or absence of “key individuals” in the group ^77^. Such key individuals are known to catalyse intergroup violence, and the concentration of leadership towards key individuals may restrict the influence that others have and result in more unshared decisions ^77^ (low Ω).

Sharing or decentralisation of collective decision-making can promote more peaceful intergroup interactions without the presence or requirement for complex human institutions. This is not to say that institutions cannot or do not play a crucial role, but they are not a prerequisite for democratic peace to emerge in biological systems. The only necessary components are mechanisms of collective decision making and differential incentives that are intrinsic properties of humans and many other social organisms, from microbes to primates.

## ACKNOWLEDGEMENTS

We would like to thank the University of Exeter’s Behavioural Discussion Group and Mike Cant’s Socialis Lab for insightful discussions. We would also like to thank Liz Greenyer for useful discussion and Hattie Lavender for proof-reading.

## Authors contributions

Conceptualisation and writing the manuscript: KLH and DWES, with input from all co-authors and extensive support from MAC. Analysis and implementation: DWES and KLH, with input from MP.

## Data and materials availability

Scripts are available to download from https://github.com/sankeydan/demoPeace2/

## SUPPLEMENTAL MATERIAL

### Materials and Methods

#### Running the model

We initialise a model with the probability of Hawk playing for both leader and follower classes at (*P*_*L*_ *= P*_*F*_ *=* 0.01). This initial value does not change the model outcome. Using these values, and chosen values for Ω, ε and *N* (although see Fig. S1 for how *N* makes no qualitative difference to results) we first calculate *P, C*_*L*_, *C*_*F*_, *V*_*L*_ and *V*_*F*_ using Equations 1-5 respectively. These values are then used to calculate the fitness of leader strategies Hawk, *W*_*LH*_, or Dove, *W*_*LD*_, and the fitness of follower strategies Hawk, *WFH*, or Dove, *W*_*FD*_, using Equations 6-9 respectively. If Hawk provides a larger fitness payoff than Dove then the probability of playing Hawk will increase for that class, and vice versa if Dove performs better, the probability of playing Hawk decreases for that class. The amount that the probability of Hawk-playing increases or decreases by in each round is not constant, it decays exponentially in each round, *r*, governed by the function 0.7e^(−0.08 * *r*)^, such that in the first round the changes are large (0.65) but by the 300^th^ round the increase for successful Hawks or decrease for successful Doves is negligible (2 * 10^−11^). It may seem that we have varied the strength of selection, however this is purely an endeavour of algorithm optimisation. We have added or subtracted incremental and equal changes to strategies of 0.001 in each round (same strength of selection in each round) and find the same results (Fig. S5). The exact values of 0.7 and 0.08 can be chosen arbitrarily; though these present values have the result that classes converge in a small number of rounds. Strategies have converged even by the 100^th^ round (Fig. S5), though we run the model to the 300^th^ round to ensure precise results.

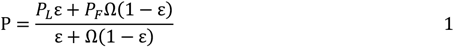

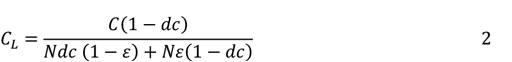

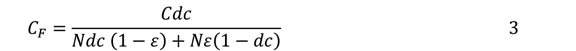

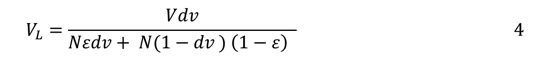

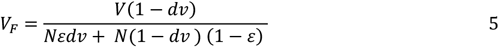

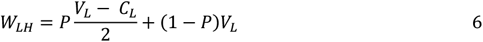

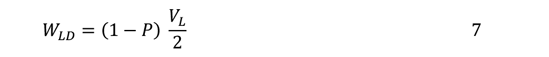

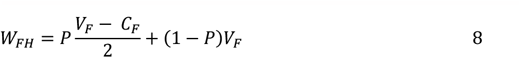

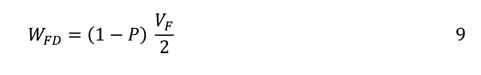

**Figure S1:**
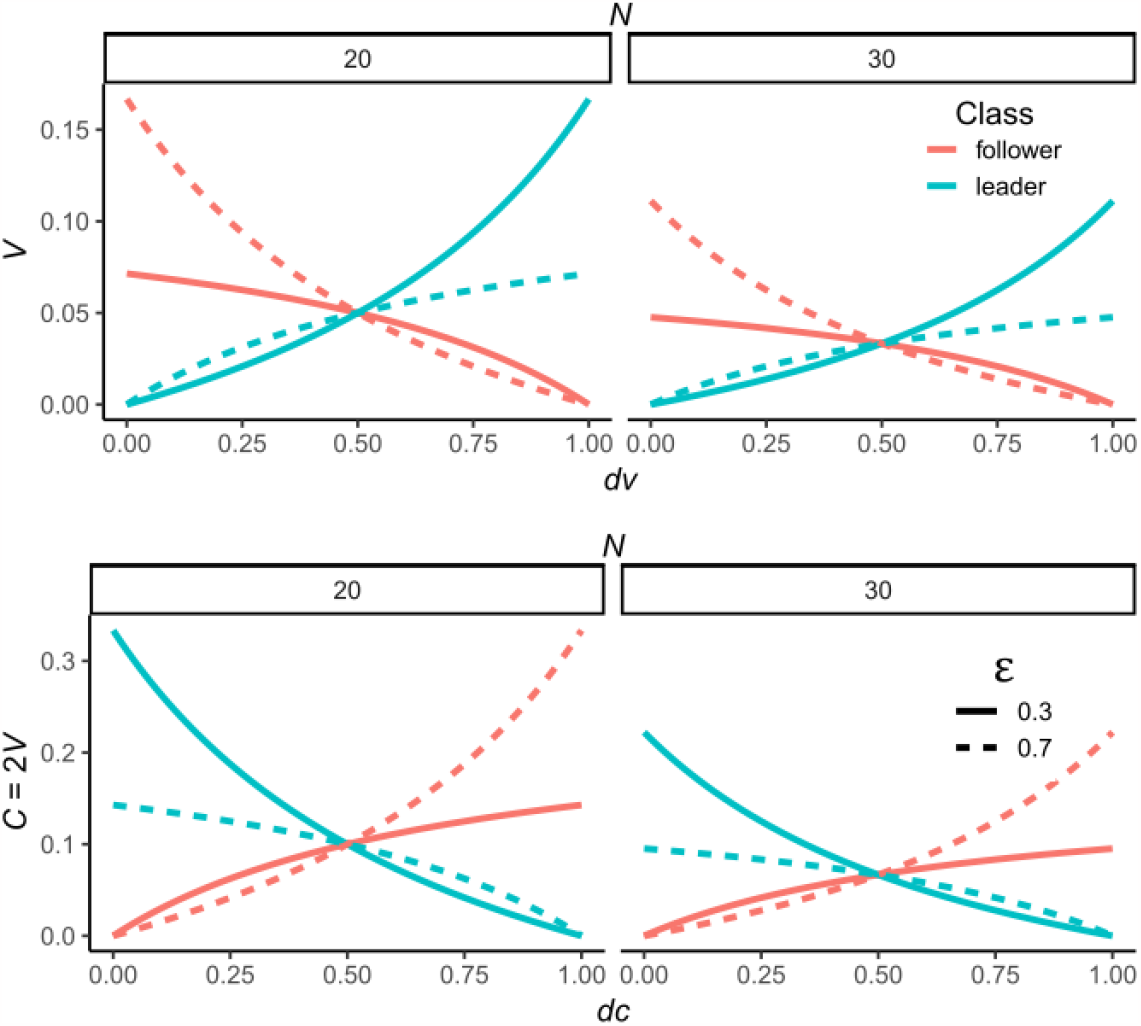
The share of benefits *(top row*) and costs *(bottom row*) obtained by each individual leader (*blue*) and follower (*pink*) under different values of *dv* and *dc* and *N*. Shares of costs and benefits are calculated using Equations 2-5 described in Materials and Methods. When *dv* = 0.5 and *dc* = 0.5, leader’s and follower’s receive identical shares, whereas values above 0.5 describe division rules with “advantaged” leaders who pay less of the costs or gain more of the benefits from fighting than followers, and vice versa followers are advantaged for values below 0.5. Note that the impact of changing group size (*N*) from 20 (*left*) to 30 (*right*) changes the absolute values of each share by rescaling the y-axis but importantly does not change the proportional relationship describing what each leader and follower receives relative to one another. Thus, *N* does not change any of the presented results. Parameter values: *C* = 2*V, dc* = 0.5 (top row) and *dv* = 0.5 (bottom row).

**Figure S2:**
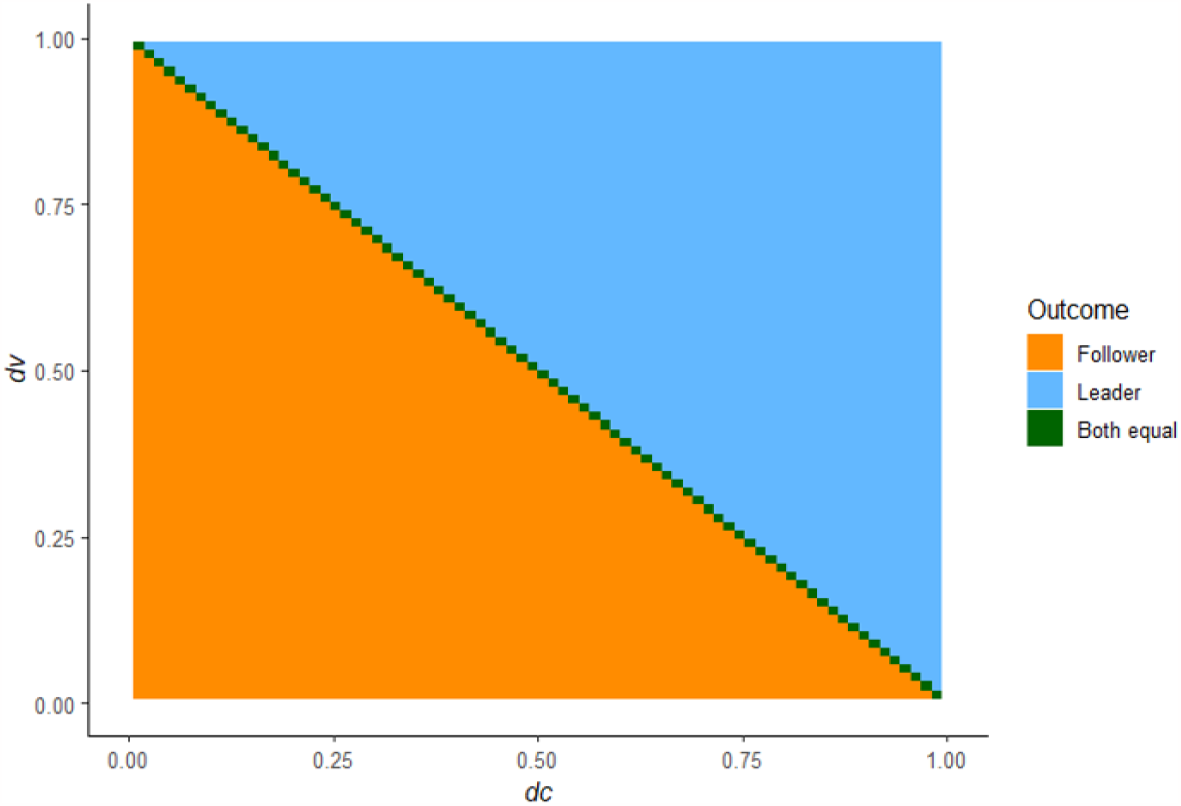
Advantaged class have higher optimal Hawk-playing probabilities. Leaders prefer more aggressive interactions by playing Hawk more often than followers (*V*_*L*_ / *C*_*L*_ > *V*_*F*_ / *C*_*F*_) when 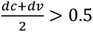(shown in blue). Followers have a higher preference than leaders (*V*_*L*_ / *C*_*L*_ > *V*_*F*_ / *C*_*F*_) when 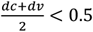(shown in orange). Leaders and followers have an equal preference (*V*_*F*_ / *C*_*F*_ =*V*_*L*_ / *C*_*L*_) when 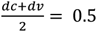 (shown in green). We use the terms *advantaged* and *disadvantaged* to describe the class which gains more from interactions with outgroups. For example, leaders are described as *advantaged* or *disadvantaged* when the combined sharing rules of *dc* and *dv* are to the right or left of the diagonal green line respectively.

**Figure S3:**
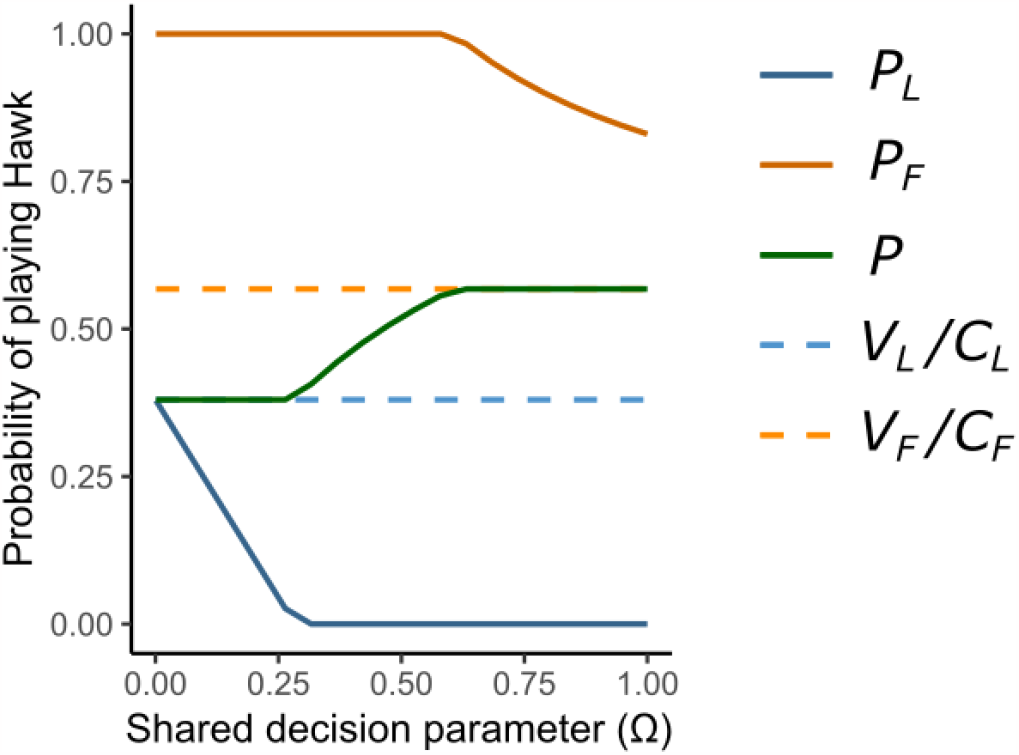
Illustrative example of the potential for “democratic war”. The propensity to play Hawk for leaders (*P*_*L*_) and followers (*P*_*F*_) and the probability of playing Hawk across the population (*P*) are shown across shared decision parameter (Ω). The population strategy rapidly transitions from leaders’ preference (*V*_*L*_ / C_L_) to followers’ preference (*V*_*F*_ / *C*_*F*_). Note that the followers have a higher optimum preference than the leaders because they are advantaged in this example, i.e., 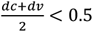.Parameter values: ε = 0.3, *C* = 2*V, dc* = 0.45, *dv* = 0.45.

**Figure S4.**
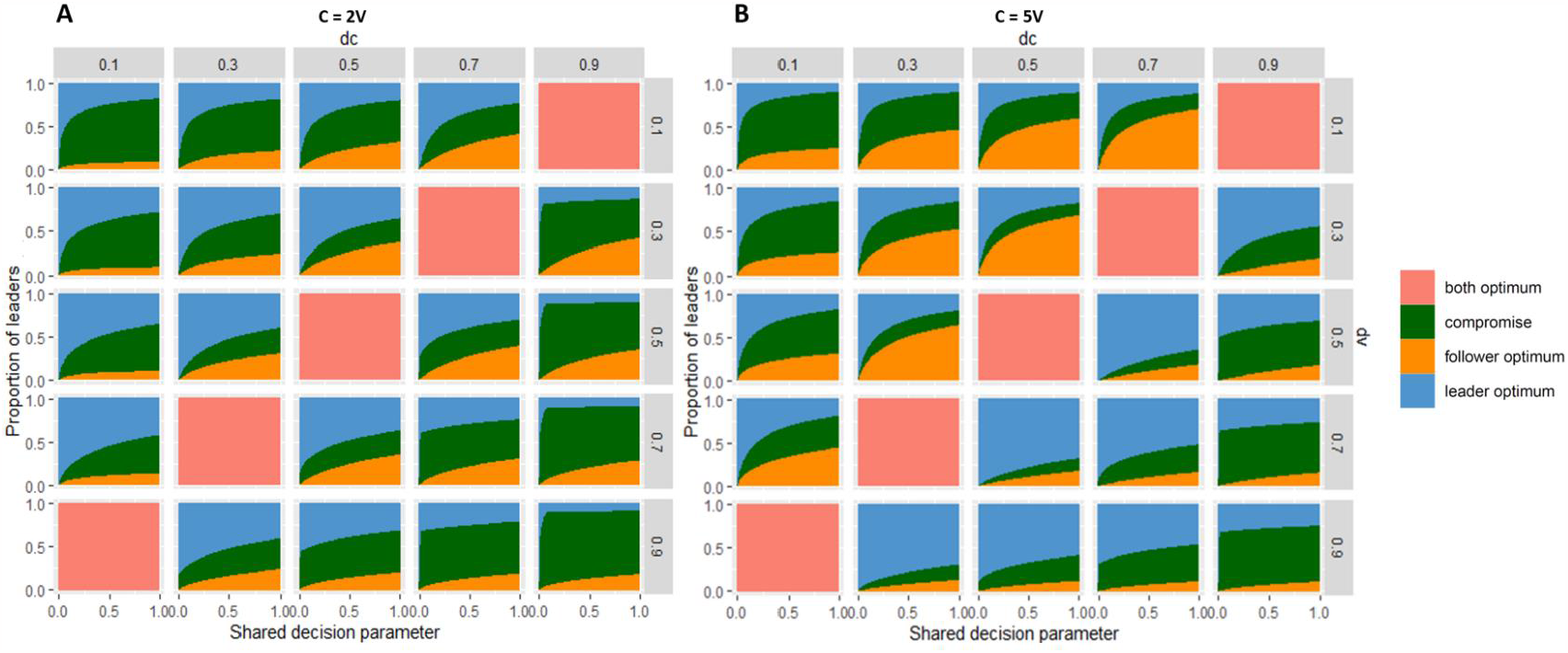
Group’s evolved strategy in response to Ω and ε also depends on *dv, dc* and *C*:*V* ratio. Related to Figure 3. Population’s evolved strategy as either the follower’s optimum preference (*P* = *V*_*F*_ / *C*_*F*_, shown in orange), leader’s optimum preference (*P* = *V*_*L*_ / *C*_*L*_, shown in blue), in compromise (*P* is between *V*_*F*_ / *C*_*F*_ and *V*_*L*_ / *C*_*L*_, shown in green) or both classes satisfying optimum preferences (*P* = *V*_*L*_ / *C*_*L*_ = *V*_*F*_ / *C*_*F*_, shown in salmon). The share of costs, *dc*, and benefits, *dv* – in which increasing values favour leaders – vary between plots on the x and y axis respectively. Regardless of *dc* and *dv*, when leaders monopolise decision (Ω is low) and when leaders are relatively abundant (ε is high) the group’s evolved strategy is likely to be at the leader’s optimum. When the mean of *dv* and *dc* are closer to 0 or 1, we see more compromise states (green). This is because there is a greater difference between each classes’ optimal levels of Hawk-playing, with one classes’ strategy close to obligate Dove and the others’ close to obligate Hawk. This minimises the anchoring ability of both parties to sway the collective decision in their favour, leading to more compromise outcomes. For more on anchoring see section “loudest voice prevails” from the Main Text. When the mean of *dv* and *dc* = 0.5 each classes’ optima are equal (*V*_*L*_ / *C*_*L*_ = *V*_*F*_ / *C*_*F*_) meaning that both classes will have the same strategy (salmon). Cost of fighting *C* = 2*V* in **A** and *C* = 5*V* in **B**. The increased cost of fighting decreases optimal Hawk playing for both parties. It is thus the class which is disadvantaged which lose anchoring ability. The result is clearly illustrated by the data to the left and right of the salmon data in **B**. To the left, when the followers are advantaged, they have more ability to sway the decision, but leaders have more sway to the right when they are advantaged. This trend is absent in **A** where costs are lower. The higher the costs, the more a class with inherent advantage can turn their higher aggressiveness into influence, though overall strategies are more peaceful.

**Figure S5.**
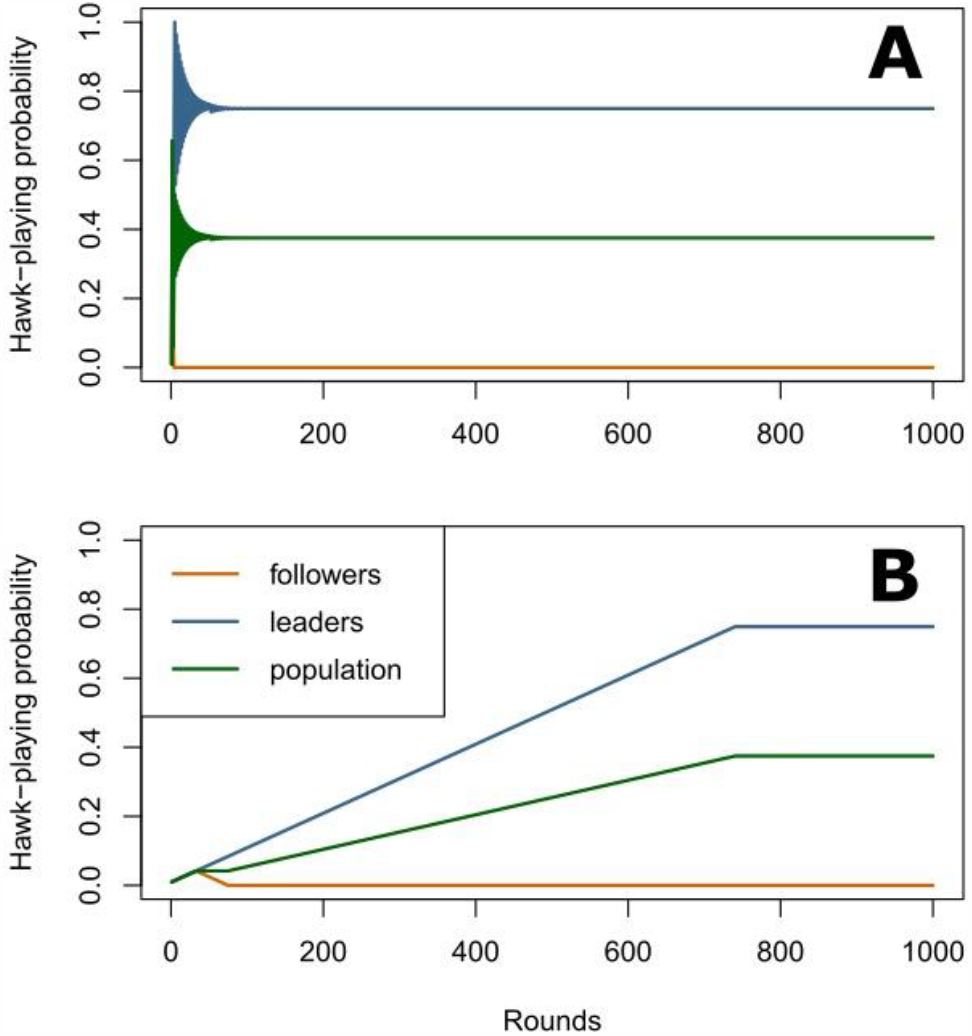
Algorithm optimisation. **A)** Method used to generate main results involves a successive decrease in the magnitude used to update the strategy in each progressive round. This allows us to reach the evolutionarily stable strategy faster (N=300 rounds used in main results) than the method in **B)** where strategy differences are always 0.001 in each round. The advantage of our method is savings in computational time and costs, despite obtaining the same end result. Parameter values: *C* = 8*V, dv* = 0.75, *dc* = 0.75, Ω = 1, ε = 0.5.

## REFERENCES

1. Choi, J.-K., and Bowles, S. (2007). The coevolution of parochial altruism and war. Science (80-.). 318, 636–640.

2. Gat, A. (2008). War in Human Civilisation (Oxford University Press).

3. Keeley, L.H. (1996). War Before Civilization: the Myth of the Peaceful Savage (Oxford University Press).

4. Rummel, R.J. (1997). Death by government (Transaction Publishers).

5. Allen, M.W., Bettinger, R.L., Codding, B.F., Jones, T.L., and Schwitalla, A.W. (2016).Resource scarcity drives lethal aggression among prehistoric hunter-gatherers in central California. Proc. Natl. Acad. Sci. 113, 12120–12125.

6. Lahr, M.M., Rivera, F., Power, R.K., Mounier, A., Copsey, B., Crivellaro, F., Edung, J.E., Fernandez, J.M., Kiarie, C., and Lawrence, J. (2016). Inter-group violence among early Holocene hunter-gatherers of West Turkana, Kenya. Nature 529, 394–398.

7. Buzzell, E., and Preston, S.H. (2007). Mortality of American troops in the Iraq war. Popul. Dev. Rev. 33, 555–566.

8. Fearon, J.D. (1995). Rationalist Explanations for War. Int. Organ. 49, 379–414.

9. Hames, R. (2020). Cultural and reproductive success and the causes of war: A Yanomamö perspective. Evol. Hum. Behav. 41, 183–187.

10. Durrant, R. (2011). Collective violence: An evolutionary perspective. Aggress. Violent Behav. 16, 428–436.

11. Glowacki, L., Wilson, M.L., and Wrangham, R.W. (2020). The evolutionary anthropology of war. J. Econ. Behav. Organ. 178, 963–982.

12. Goncalves, I.B., Morris-Drake, A., Kennedy, P., and Radford, A.N. (2022). Fitness consequences of outgroup conflict. Elife 11, e74550.

13. Johnstone, R.A., Cant, M.A., Cram, D., and Thompson, F.J. (2020). Exploitative leaders incite intergroup warfare in a social mammal. Proc. Natl. Acad. Sci. 117, 29759–29766.

14. Wrangham, R.W., Wilson, M.L., and Muller, M.N. (2006). Comparative rates of violence in chimpanzees and humans. Primates 47, 14–26.

15. Kant, I. (2003). To perpetual peace: A philosophical sketch (T. Humphrey, Trans)(1795). Indianap. Hackett.

16. Paine, T. (2008). Rights of Man, Common Sense, and other political writings (Oxford University Press).

17. Tocqueville, A. (2003). Democracy in America: And two essays on America (Penguin UK).

18. Mello, P.A. (2014). Democratic Peace Theory From Kant’s “Perpetual Peace” to Democratic Peace Theoretical Explanations for the Democratic Peace. SAGE Encycl. War Soc. Sci. Perspect.

19. Ray, J.L. (1995). Democracy and international conflict: An evaluation of the democratic peace proposition (University of South Carolina Press).

20. Sankey, D.W.E., Hunt, K.L., Croft, D.P., Franks, D.W., Green, P.A., Thompson, F.J., Johnstone, R.A., and Cant, M.A. (2022). Leaders of war: modelling the evolution of conflict among heterogeneous groups. Philos. Trans. R. Soc. B 377, 20210140.

21. Rummel, R.J. (1995). Democracies are less warlike than other regimes. Eur. J. Int. relations 1, 457–479.

22. Benoit, K. (1996). Democracies really are more pacific (in general) reexamining regime type and war involvement. J. Conflict Resolut. 40, 636–657.

23. Dafoe, A., Oneal, J.R., and Russett, B. (2013). The democratic peace: Weighing the evidence and cautious inference. Int. Stud. Q. 57, 201–214.

24. Imai, K., and Lo, J. (2021). Robustness of empirical evidence for the democratic peace: A nonparametric sensitivity analysis. Int. Organ. 75, 901–919.

25. Russett, B.M., and Oneal, J.R. (2001). Triangulating peace: Democracy,interdependence, and international organizations (WW Norton & Company Incorporated).

26. Gartzke, E. (2007). The capitalist peace. Am. J. Pol. Sci. 51, 166–191.

27. Simpson, S. (2019). Making liberal use of Kant? Democratic peace theory and Perpetual Peace. Int. Relations 33, 109–128.

28. Smith, J.E., Estrada, J.R., Richards, H.R., Dawes, S.E., Mitsos, K., and Holekamp, K.E. (2015). Collective movements, leadership and consensus costs at reunions in spotted hyaenas. Anim. Behav. 105, 187–200.

29. Smith, J.E., Gavrilets, S., Mulder, M.B., Hooper, P.L., Mouden, C. El Nettle, D., Hauert, C., Hill, K., Perry, S., Pusey, A.E., et al. (2016). Leadership in Mammalian Societies: Emergence, Distribution, Power, and Payoff. Trends Ecol. Evol. 31, 54–66.

30. King, A.J., Douglas, C.M.S., Huchard, E., Isaac, N.J.B., and Cowlishaw, G. (2008). Dominance and affiliation mediate despotism in a social primate. Curr. Biol. 18, 1833–1838.

31. Lusseau, D., and Conradt, L. (2009). The emergence of unshared consensus decisions in bottlenose dolphins. Behav. Ecol. Sociobiol. 63, 1067–1077.

32. McComb, K., Shannon, G., Durant, S.M., Sayialel, K., Slotow, R., Poole, J.H., and Moss, C.J. (2011). Leadership in elephants: the adaptive value of age. Proc. R. Soc. London B Biol. Sci. 278, 3270–3276.

33. Brent, L.J.N., Franks, D.W., Foster, E.A., Balcomb, K.C., Cant, M.A., and Croft, D.P. (2015). Ecological knowledge, leadership, and the evolution of menopause in killer whales. Curr. Biol. 25, 746–750.

34. Strandburg-Peshkin, A., Clutton-Brock, T., and Manser, M.B. (2020). Burrow usage patterns and decision-making in meerkat groups. Behav. Ecol. 31, 292–302.

35. Strandburg-Peshkin, A., Farine, D.R., Couzin, I.D., and Crofoot, M.C. (2015). Shared decision-making drives collective movement in wild baboons. Science (80-.). 348, 1358–1361.

36. Papageorgiou, D., and Farine, D.R. (2020). Shared decision-making allows subordinates to lead when dominants monopolize resources. Sci. Adv. 6, eaba5881.

37. Yomosa, M., Mizuguchi, T., Vásárhelyi, G., and Nagy, M. (2015). Coordinated behaviour in pigeon flocks. PLoS One 10, 1–17.

38. Whitehead, H. (2016). Consensus movements by groups of sperm whales. Mar. Mammal Sci. 32, 1402–1415.

39. Gall, G.E.C., Strandburg-Peshkin, A., Clutton-Brock, T.H., and Manser, M.B. (2017). As dusk falls: collective decisions about the return to sleeping sites in meerkats. Anim. Behav. 132, 91–99.

40. Walker, R.H., King, A.J., McNutt, J.W., and Jordan, N.R. (2017). Sneeze to leave: African wild dogs (Lycaon pictus) use variable quorum thresholds facilitated by sneezes in collective decisions. Proc. R. Soc. B Biol. Sci. 284.

41. Furuichi, T. (2020). Variation in intergroup relationships among species and among and within local populations of African apes. Int. J. Primatol. 41, 203–223.

42. Popat, R., Cornforth, D.M., McNally, L., and Brown, S.P. (2015). Collective sensing and collective responses in quorum-sensing bacteria. J. R. Soc. Interface 12, 20140882.

43. Granato, E.T., Meiller-Legrand, T.A., and Foster, K.R. (2019). The evolution and ecology of bacterial warfare. Curr. Biol. 29, R521–R537.

44. Tokuyama, N., and Furuichi, T. (2017). Leadership of old females in collective departures in wild bonobos (Pan paniscus) at Wamba. Behav. Ecol. Sociobiol. 71, 1–10.

45. Smith, J.M., and Price, G.R. (1973). The logic of animal conflict. Nature 246, 15–18.

46. Mathew, S., and Boyd, R. (2011). Punishment sustains large-scale cooperation in prestate warfare. Proc. Natl. Acad. Sci. U. S. A. 108, 11375–11380.

47. Arseneau-Robar, T.J.M., Taucher, A.L., Müller, E., van Schaik, C., Bshary, R., and Willems, E.P. (2016). Female monkeys use both the carrot and the stick to promote male participation in intergroup fights. Proc. R. Soc. B Biol. Sci. 283, 20161817.

48. Kingma, S.A., Santema, P., Taborsky, M., and Komdeur, J. (2014). Group augmentation and the evolution of cooperation. Trends Ecol. Evol. 29, 476–484.

49. Sääksvuori, L. (2014). Intergroup conflict, ostracism, and the evolution of cooperation under free migration. Behav. Ecol. Sociobiol. 68, 1311–1319.

50. Gavrilets, S., Auerbach, J., and Van Vugt, M. (2016). Convergence to consensus in heterogeneous groups and the emergence of informal leadership. Sci. Rep. 6, 1–10.

51. Pyritz, L.W., King, A.J., Sueur, C., and Fichtel, C. (2011). Reaching a consensus: Terminology and concepts used in coordination and decision-making research. Int. J. Primatol. 32, 1268–1278.

52. Glowacki, L., and Wrangham, R. (2015). Warfare and reproductive success in a tribal population. Proc. Natl. Acad. Sci. U. S. A. 112, 348–353.

53. Gavrilets, S., and Fortunato, L. (2014). A solution to the collective action problem in between-group conflict with within-group inequality. Nat. Commun. 5, 1–11.

54. Ioannou, C.C., Rocque, F., Herbert-Read, J.E., Duffield, C., and Firth, J.A. (2019). Predators attacking virtual prey reveal the costs and benefits of leadership. Proc. Natl. Acad. Sci. 116, 8925–8930.

55. Johnstone, R.A., and Manica, A. (2011). Evolution of personality differences in leadership. Proc. Natl. Acad. Sci. U. S. A. 108, 8373–8.

56. Rands, S.A., Cowlishaw, G., Pettifor, R.A., Rowcliffe, J.M., and Johnstone, R.A. (2003). Spontaneous emergence of leaders and followers in foraging pairs. Nature 423, 432–434.

57. Heinsohn, R., and Packer, C. (1995). Complex Cooperative Strategies in Group-Territorial African Lion. Science (80-.). 269, 1260–1263.

58. Haig, D. (1997). Parental antagonism, relatedness asymmetries, and genomic imprinting. Proc. R. Soc. London. Ser. B Biol. Sci. 264, 1657–1662.

59. Haig, D. (1996). Placental hormones, genomic imprinting, and maternal—fetal communication. J. Evol. Biol. 9, 357–380.

60. Pamilo, P., and Rosengren, R. (1983). Sex ratio strategies in Formica ants. Oikos, 24–35.

61. Boomsma, J.J., and Grafen, A. (1991). Colony-level sex ratio selection in the eusocial Hymenoptera. J. Evol. Biol. 4, 383–407.

62. King, D.C., and Zeckhauser, R.J. (2002). Punching and counter-punching in the US Congress: Why party leaders tend to be extremists. In Conference on Leadership.

63. Davis, G.H., Crofoot, M.C., and Farine, D.R. (2022). Using optimal foraging theory to infer how groups make collective decisions. Trends Ecol. Evol.

64. Lee, H.C., and Teichroeb, J.A. (2016). Partially shared consensus decision making and distributed leadership in vervet monkeys: older females lead the group to forage. Am. J. Phys. Anthropol. 161, 580–590.

65. Stanford, C.B. (1998). The social behavior of chimpanzees and bonobos: empirical evidence and shifting assumptions. Curr. Anthropol. 39, 399–420.

66. Wrangham, R.W., and Glowacki, L. (2012). Intergroup aggression in chimpanzees and war in nomadic hunter-gatherers. Hum. Nat. 23, 5–29.

67. Goodall, J. (1986). The chimpanzees of Gombe: Patterns of behavior. Cambridge Mass.

68. Smith, J.E., Fichtel, C., Holmes, R.K., Kappeler, P.M., van Vugt, M., and Jaeggi, A. V (2022). Sex bias in intergroup conflict and collective movements among social mammals: male warriors and female guides. Philos. Trans. R. Soc. B 377, 20210142.

69. Smith, J.E., Ortiz, C.A., Buhbe, M.T., and van Vugt, M. (2020). Obstacles and opportunities for female leadership in mammalian societies: A comparative perspective. Leadersh. Q. 31, 101267.

70. Ceccarelli, E., Rangel Negrín, A., Coyohua-Fuentes, A., Canales-Espinosa, D., and Dias, P.A.D. (2020). Sex differences in leadership during group movement in mantled howler monkeys (Alouatta palliata). Am. J. Primatol. 82, e23099.

71. Tecot, S.R., and Romine, N.K. (2012). Leading Ladies: Leadership of Group Movements in a Pair-Living, Co-Dominant, Monomorphic Primate Across Reproductive Stages and Fruit Availability Seasons. Am. J. Primatol. 74, 591–601.

72. Kurvers, R.H.J.M., Eijkelenkamp, B., van Oers, K., van Lith, B., van Wieren, S.E., Ydenberg, R.C., and Prins, H.H.T. (2009). Personality differences explain leadership in barnacle geese. Anim. Behav. 78, 447–453.

73. Nakayama, S., Harcourt, J.L., and Johnstone, R.A. (2012). Initiative, Personality and Leadership in Pairs of Foraging Fish. 7.

74. Sasaki, T., Mann, R.P., Warren, K., Herbert, T., Wilson, T., and Biro, D. (2017). Personality and the collective: Bold homing pigeons occupy higher leadership ranks in flocks. Philos. Trans. R. Soc. B Biol. Sci.

75. Gilby, I.C., Eberly, L.E., and Wrangham, R.W. (2008). Economic profitability of social predation among wild chimpanzees: individual variation promotes cooperation. Anim. Behav. 75, 351–360.

76. Gilby, I.C., Machanda, Z.P., Mjungu, D.C., Rosen, J., Muller, M.N., Pusey, A.E., and Wrangham, R.W. (2015). ‘Impact hunters’ catalyse cooperative hunting in two wild chimpanzee communities. Philos. Trans. R. Soc. B Biol. Sci. 370, 20150005.

77. Glowacki, L., and McDermott, R. (2022). Key individuals catalyse intergroup violence. Philos. Trans. R. Soc. B 377, 20210141.

78. Massaro, A.P., Gilby, I.C., Desai, N., Weiss, A., Feldblum, J.T., Pusey, A.E., and Wilson, M.L. (2022). Correlates of individual participation in boundary patrols by male chimpanzees. Philos. Trans. R. Soc. B 377, 20210151.

79. Arseneau-Robar, T.J.M., Taucher, A.L., Schnider, A.B., van Schaik, C.P., and Willems, E.P. (2017). Intra- and interindividual differences in the costs and benefits of intergroup aggression in female vervet monkeys. Anim. Behav. 123, 129–137.

80. Riehl, C., and Frederickson, M.E. (2016). Cheating and punishment in cooperative animal societies. Philos. Trans. R. Soc. B Biol. Sci. 371, 20150090.

